# Phase separation of mycobacterial Rho factor is associated with acid stress

**DOI:** 10.1101/2025.08.28.672908

**Authors:** Sofia M. Moreira, Joseph T. Wade, Chris M. Brown

## Abstract

Rho is a bacterial transcription termination factor that contains additional regions in some species, including many mycobacteria. These additional regions are compositionally biased and are predicted to be intrinsically disordered and associated with Liquid-Liquid Phase Separation (LLPS). However, LLPS has only been experimentally demonstrated for Rho from *Bacteroides thetaiotaomicron*. Here, we extend a prior bioinformatic analysis of Rho from mycobacterial species, where Rho has additional regions that differ from Rho in *B. thetaiotaomicron*. We show that *Mycolicibacterium smegmatis* Rho forms *in vitro* and *in vivo* condensates by LLPS, and that LLPS depends upon the additional regions. *In vitro*, droplet formation for Rho from four mycobacterial species was modulated by the presence of RNA and was observed for the intrinsically disordered regions alone, suggesting that these regions are sufficient for droplet formation. Moreover, Rho phase separation in living *M. smegmatis* cells occurs during acid stress and may impact cell fitness. This phenotype is due to Rho’s additional regions, since engineered mycobacterial Rho lacking the additional regions does not undergo LLPS. In summary, our results indicate that LLPS is a widespread mechanism in which mycobacterial Rho assembles into condensates, possibly terminating the transcription of different genes, which allows tolerance to stress conditions.

## 1.1 INTRODUCTION

Widespread in eukaryotic cells, membraneless organelles are isolated compartments where frequent exchanges of substances occur (Hirose et al. 2023). One mechanism suggested for the assembly of such structures is liquid-liquid phase separation (LLPS) in which the biomolecules separate in solution, creating a dense liquid phase (resembling liquid droplets) and a dilute surrounding phase (Alberti et al. 2019). Usually, proteins that can undergo LLPS are composed of intrinsically disordered regions (IDR) and/or repeats of interaction domains that confer the ability to phase-separate (De Sancho 2022). Numerous eukaryotic cellular processes rely on the proper regulation of LLPS, and severe human diseases are associated with anomalies in LLPS (Boyko and Surewicz 2022; Chen et al. 2022; Mehta and Zhang 2022).

Although the molecular mechanism of LLPS is better characterized in eukaryotes, recent studies also demonstrated this phenomenon in bacterial cells (Azaldegui et al. 2021; Guo et al. 2024). The organization of biochemical reactions in membraneless organelles allows bacteria to regulate the activity of proteins (Guilhas et al. 2020; Heinkel et al. 2019; Ladouceur et al. 2020; Lasker et al. 2020; Monterroso et al. 2019; Sanchez et al. 2015; Saurabh et al. 2022; Tan et al. 2022; Wang et al. 2019) and increase fitness during stress conditions (Al-Husini et al. 2018; Goldberger et al. 2022; Harami et al. 2020; Janissen et al. 2018; Jin et al. 2021; Krypotou et al. 2023; Pu et al. 2019; Zhao et al. 2019). For example, the regulation of asymmetric cell division in *Caulobacter crescentus* is associated with the phase separation of PodJ (Tan et al. 2022). In this same species, LLPS was also demonstrated to modulate the activity of PopZ by concentrating and coordinating kinase signaling (Lasker et al. 2020; Saurabh et al. 2022) as well as of RNP bodies by facilitating mRNA decay (Al-Husini et al. 2018). During harsh conditions (e.g., high salt and nutrient starvation), Hfq forms polar condensates with RNA in *Escherichia coli* that are essential for its posttranscriptional roles as an RNA chaperone (Goldberger et al. 2022). In *Bacteroides thetaiotaomicron*, survival during carbon deprivation in the mouse gut is associated with phase separation of the Rho protein (_Bth_Rho). Phase separation of _Bth_Rho is promoted by an IDR (Krypotou et al. 2023).

Rho is a hexameric RNA-dependent ATPase responsible for factor-dependent transcription termination in bacteria (Mitra et al. 2017). While absent in Cyanobacteriota, Mycoplasmatota, and a few members of Bacillota, Rho is widespread and well-conserved among bacterial species (D’Heygère et al. 2013; Moreira et al. 2024). Rho typically has three main domains: N-terminal, RNA-binding, and Rho ATPase. However, some species have atypical Rho proteins that contain at least one “additional” region with distinct properties (Krypotou et al. 2023; Mitra et al. 2014; Moreira et al. 2024; Nowatzke et al. 1997; Pallarès et al. 2016; Simon et al. 2021; Trzilova et al. 2020; Yuan and Hochschild 2017). Notably, the atypical Rho of *Clostridium botulinum* harbors an internal prion-like region that self-assembles into structures similar to amyloids that alter the conformation of this protein based on environmental changes (Pallarès et al. 2016; Yuan and Hochschild 2017). *In vitro* transcription termination experiments have shown that some Rho insertions are involved with increasing the affinity for RNA in *B. fragilis* (Simon et al. 2021), *Micrococcus luteus* (Nowatzke et al. 1997), and *Mycobacterium tuberculosis* (Mitra et al. 2014). These insertions have compositionally biased regions containing amino acids commonly associated with phase separation (arginine, asparagine, aspartic acid, and glycine) (Azaldegui et al. 2021). However, there is currently only one study reporting *in vivo* and *in vitro* LLPS of Rho, that of _Bth_Rho (Krypotou et al. 2023).

Rho is essential in many bacterial species, and its gene inactivation causes the rapid death of *M. tuberculosis* (Botella et al. 2017). *M. tuberculosis* Rho (_Mtb_Rho) is an atypical Rho that has 602 amino acids, including two additional regions: an initial region of 36 amino acids before the N-terminal domain, and an insertion region of 152 amino acids between the N-terminal and RNA-binding domains (Figure 1A). The additional regions of _Mtb_Rho are predicted to form an IDR and were not resolved in the cryo-EM structure of the _Mtb_Rho hexamer (Saridakis et al. 2022). Since proteins with IDRs are associated with LLPS, we speculated that _Mtb_Rho forms condensates under certain physiological conditions. To date, LLPS in *M. tuberculosis* has been proposed for only one protein, the ABC transporter Rv1747 (Heinkel et al. 2019).

**Figure 1:**
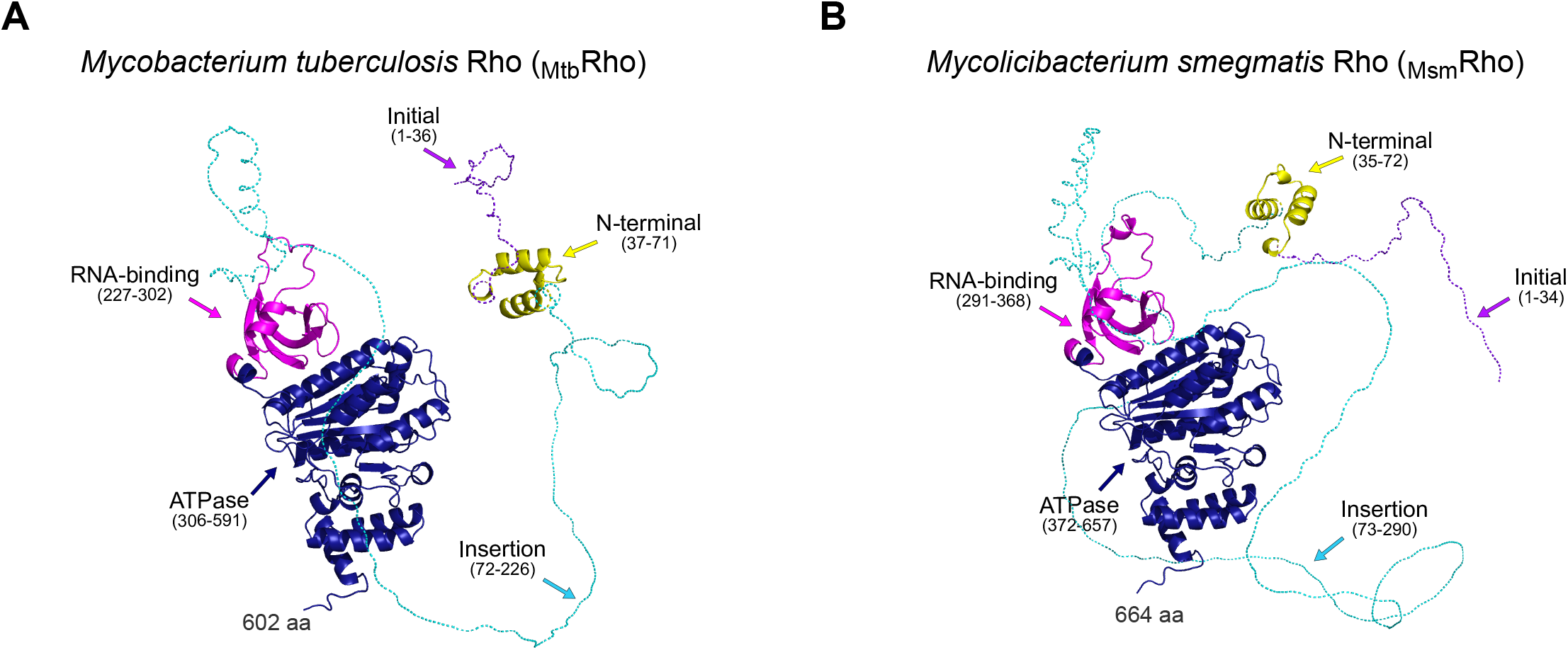
Monomer of the transcription factor Rho in two important mycobacterial species. (A) *M. tuberculosis* Rho (_Mtb_Rho) (AlphaFold ID: AF-P9WHF3-F1). (B) *M. smegmatis Rho* (_Msm_Rho) (AlphaFold ID: AF-I7G6F1-F1). 3D structures were obtained from the AlphaFold database and were colored according to the different domains, with the additional regions represented as dashed lines.

*Mycolicibacterium smegmatis* is a model organism for studying mycobacteria (Sparks et al. 2023). A monomer of *M. smegmatis* Rho (_Msm_Rho) has 664 amino acids with an initial (34 amino acids) and an insertion (217 amino acids) region (Figure 1B), and it shares 76.21% amino acid identity with _Mtb_Rho. Inspired by the study performed by Krypotou et al. (2023), and given the importance of Rho in Mycobacteria, we tested whether _Msm_Rho undergoes phase separation *in vitro* and *in vivo*. Our results show that the additional regions of mycobacterial Rho proteins are responsible for promoting LLPS, a property that appears to be widespread across the Mycobacteriaceae family. Furthermore, we show that the ability of *M. smegmatis* Rho to phase-separate promotes resistance to acid stress.

## 2 RESULTS

### 2.1 Mycobacterial Rho have compositionally biased amino acids associated with phase separation

Mycobacteriaceae is an Actinomycetota family that includes environmental species and well-known pathogens. Previous studies of Rho in mycobacteria focused on *in vitro* transcription using *M. tuberculosis* proteins (Ahmad et al. 2023; Mitra et al. 2014; Saridakis et al. 2022). To expand the analysis of Rho in this bacterial group, we first analyzed the Rho protein sequences in 118 complete and representative mycobacterial genomes (Table S1). Almost all mycobacterial Rho factors have the three typical Rho domains (N-terminal, RNA-binding, and ATPase) and additional regions (initial and insertion) of at least 25 amino acids. The only exceptions are five species (*M. koreensis, M. insubricum, M. seoulense, M. fluoranthenivorans*, and *M. brumae*) that have an insertion region but not an initial region. In species with an initial and an insertion region, the initial region is shorter than the insertion region, with initial region length varying from 34 (*M. smegmatis* and *M. goodii*) to 80 (*M. sinensis and M. novum*) amino acids and insertion region length varying from 107 (*M. tuberculosis variant africanum*) to 268 amino acids (*M. sediminis*).

Multiple sequence alignment of mycobacterial Rho proteins showed that the typical domains are well conserved, including conservation of all expected catalytic residues (Figure S1). The additional regions are similar in species of the same genus. Among the five genera, there are some conserved sites at the beginning and the end of the initial and the insertion regions, respectively (Figure S2). Specifically, we identified four conserved sequence motifs within the initial and insertion regions (Figure 2A). The motif named “TDL” was detected at the beginning of all initial regions (Figure 2B). The three remaining motifs were detected in the insertion region and were named “RxR”, “NDQG”, and the previously reported “EDDV” (Figure 2B) (Moreira et al. 2024). Notably, there was a pattern of the distribution of these motifs that was widespread in the sequences: an isolated RxR at the center of the sequences, followed by repeats of NDQG and one copy of RxR and EDDV. This latter motif was always found immediately before the RNA-binding domain. None of these motifs was detected in the N-terminal or the Rho ATPase domain. The RNA-binding domain of mycobacteria has a small insertion in the third portion of the RNA primary binding site (RNP3) (Moreira et al. 2024), but only the NDQG motif was significantly identified in 21 of the sequences, mainly in the rapid-growing genera (*Mycobacteroides* and *Mycolicibacterium*).

**Figure 2:**
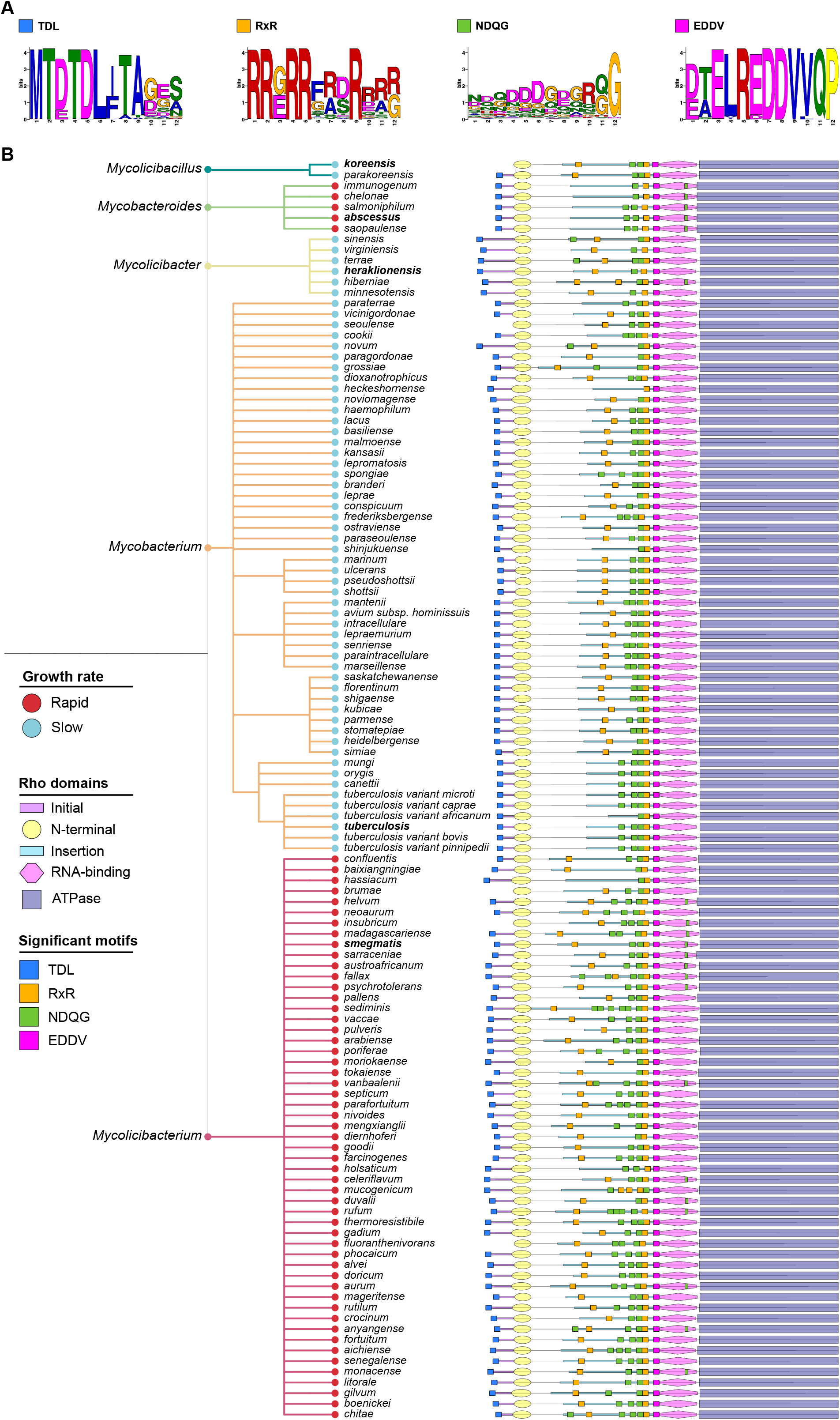
Mycobacterial Rho additional regions have significant motifs. (A) Significant motifs detected in the initial and the insertion of mycobacterial Rho factors. (B) Phylogenetic tree based on the NCBI taxonomy of 118 mycobacterial species (five genera) showing the distribution of the significant motifs along the Rho domains. Motifs were predicted by MEME and identified with MAST. Circles at the end of the nodes indicate the growth rate: rapid (red) or slow (light blue). Names in bold indicate the species which their proteins were tested for *in vitro* phase separation (Figures 3 and 4).

Besides the distribution pattern of these common motifs, there are enriched amino acids in the additional regions of mycobacterial Rho. The compositional bias is towards threonine, arginine, glycine, aspartate, and leucine. These amino acids are associated with LLPS-prone sequences (Vendruscolo and Fuxreiter 2023). In fact, the initial and insertion regions are predicted to be IDR and undergo phase separation in five representative species of the five mycobacterial genera (Figure S3). Overall, the bioinformatic analyses of mycobacterial Rho proteins show that the typical Rho domains are conserved and different from the additional regions that have distinct motifs with compositionally biased amino acids predicted to drive LLPS.

### 2.2 Mycobacterial Rho additional regions promote *in vitro* phase separation

We next evaluated the ability of Rho and the additional regions of some mycobacteria to form condensates. We performed *in vitro* LLPS experiments based on heterologous expression and purification of constructs of _m_NeonGreen-tagged _Msm_Rho (Figure 3A). We observed clear, spherical droplets of the *M. smegmatis* Rho (MsM) protein (5 µM) (Figure 3A). The positive control for our experiment was the AK390 protein (_m_NeonGreen-tagged _Bth_Rho) from Krypotou et al. (2023) that formed droplets as expected. By contrast, _m_NeonGreen alone (negative control) produced diffused fluorescence (Figure 3A). Droplets were also seen with 5 µM of the mutant IDR from _Msm_Rho that consists of the initial, N-terminal, and insertion domains (Figure 3A). Conversely, some amorphous structures, but no liquid droplets, were observed with _Msm_Rho lacking the IDR (Figure 3A). These results indicate that the IDR is necessary and sufficient for phase separation of _Msm_Rho in vitro.

**Figure 3:**
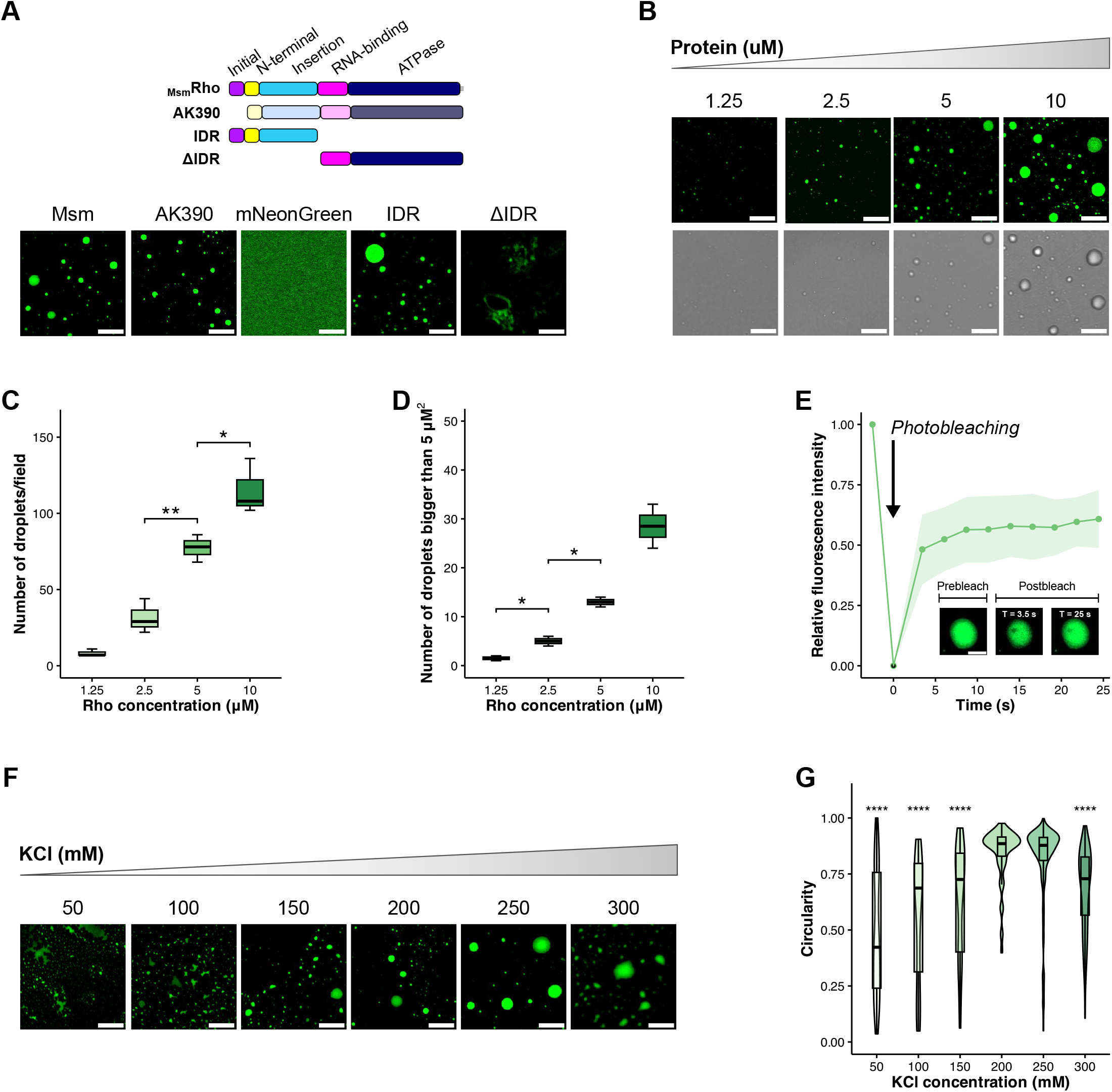
_Msm_Rho undergoes phase separation *in vitro*, which is promoted by the IDR. (A) Schematic representation of the proteins and controls used in this study (IDR, ΔIDR, _m_NeonGreen, and AK390 [_Bth_Rho from Krypotou et al. (2023)]). (B) Formation _Msm_Rho liquid droplets *in vitro*, (C) number of droplets/field, and (D) number of droplets bigger than 5 µM^2^ at increasing protein concentrations (1.25 to 10 µM). (E) Recovery curve of _Msm_Rho (n = 9 droplets) after photobleaching associated with images of one representative droplet. The fluorescence intensity of the bleached droplet was normalized to a non-bleached droplet. (F) Formation _Msm_Rho liquid droplets *in vitro* and (G) circularity of the droplets at increasing KCl concentrations (50 to 300 mM). All analyses were carried out with a buffer containing 200 mM KCl (otherwise stated) and representative images are shown. Scale bars, 5 µm; 2 µm (E). *P* values: paired t-test in (C) and (D). Kruskal-Wallis test compared to the 200 mM value (standard concentrations for the assays).

Next, we assessed factors known to be associated with *in vitro* LLPS, including protein concentration, fluorescence recovery after photobleaching (FRAP), and salt concentration (Banerjee et al. 2017; Wiedner and Giudice 2021). Consistent with LLPS, increasing concentrations of the protein (1.25 to 10 µM) (Figure 3B) led to an increase in the number of droplets (Figure 3C), and the number of large droplets (area greater than 5 µM^2^) (Figure 3D). FRAP analysis revealed that 60.8% of the fluorescence of the _m_NeonGreen tagged to _Msm_Rho could be reestablished within 25 sec, demonstrating the movement dynamics of the protein inside the droplets (Figure 3E). Increasing KCl concentration (50 to 300 mM) (Figure 3F) was associated with increased circularity of the droplets (Figure 3G); amorphous structures were observed at low (50 to 150 mM KCl) and high salt concentrations (300 mM KCl) (Figure 3G). Thus, multiple independent factors support LLPS of _Msm_Rho *in vitro*.

Many examples of LLPS involve RNA (Dignon et al. 2020). Total RNA (5 to 20 ng) of *M. smegmatis* promoted *in vitro* LLPS of _Msm_Rho (Figure 4A). Compared to the conditions without RNA, there was a significant increase in the number (Figure 4B) and size (Figure 4C) of the droplets of _Msm_Rho until the amounts of 20 ng and 5 ng, respectively. Thus, RNA appears to promote LLPS of _Msm_Rho.

**Figure 4:**
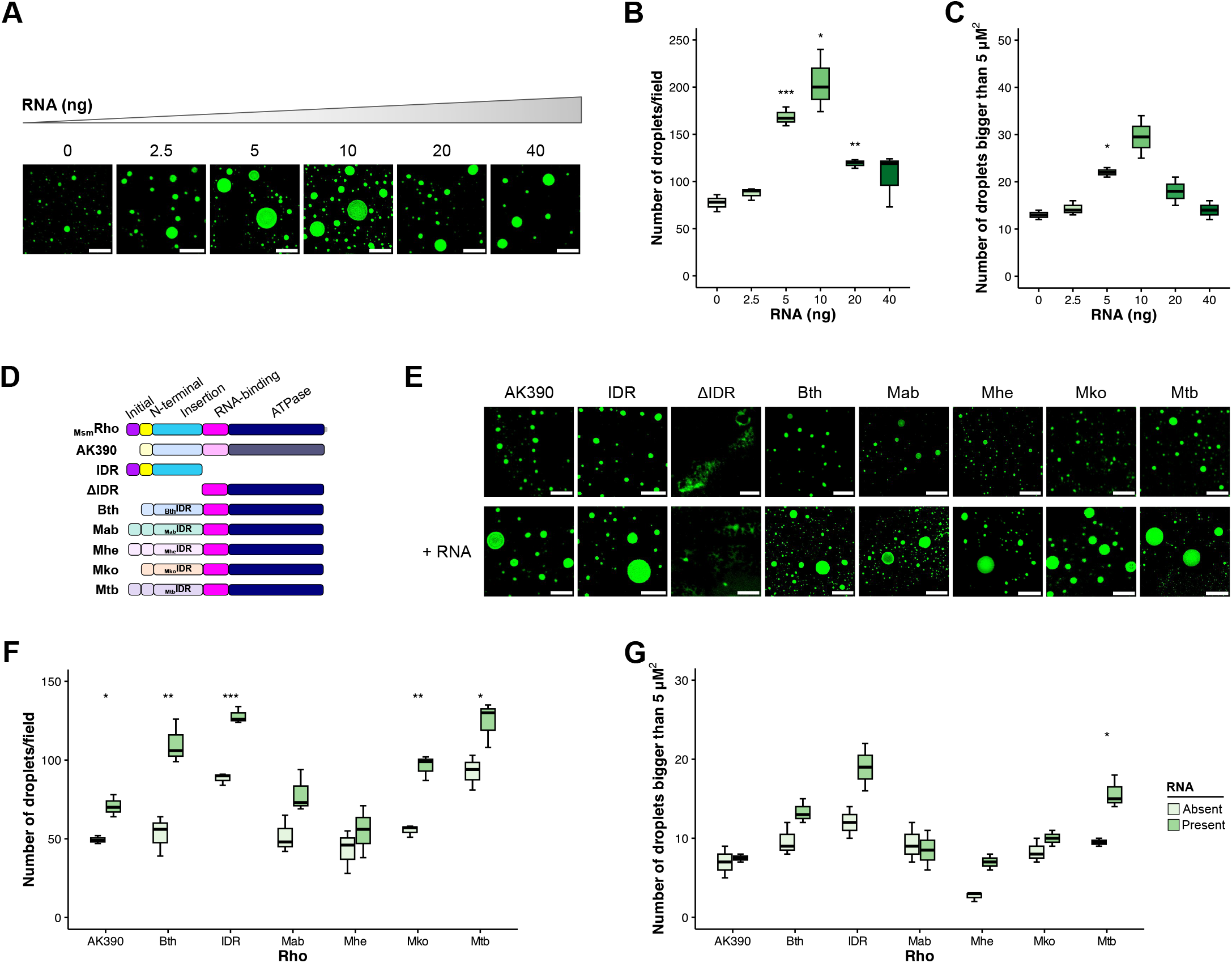
RNA promotes phase separation *in vitro* of mycobacterial Rho. (A) Formation _Msm_Rho liquid droplets *in vitro*, (B) number of droplets/field, and (C) number of droplets bigger than 5 µM^2^ at increasing amounts of the total RNA of *M. smegmatis* (2.5 to 40 µM). A) Schematic representation of the proteins used in this study. Abbreviations: Bth: *B. thetaiotaomicron*; Mab: *M. abscessus*; Mhe: *M. heraklionensis*; Mko: *M. koreensis*; Mtb: *M. tuberculosis*. (E) Formation of liquid droplets of the proteins (5 µM) represented in (D) with or without 5 ng _Msm_RNA. All analyses were carried out with a buffer containing 200 mM KCl and representative images are shown. (F) Number of droplets/field, and (G) number of droplets bigger than 5 µM^2^ with or without 5 ng RNA. Scale bars, 5 µm. *P* values: paired t-test in (B), (C), (F), (G) comparing the mean values in the absence with the presence of RNA.

Based on our bioinformatic analysis that showed conservation of the Rho typical domains and the features of mycobacterial additional regions, we created chimeric proteins containing IDRs from different mycobacterial genera and the C-terminal region from _Msm_Rho (Figure 4D). We tested IDRs from the species: *Mycobacteroides abscessus* (Mab), *Mycolicibacter heraklionensis* (Mhe), *Mycobacteroides koreensis* (Mko), and *Mycobacterium tuberculosis* (Mtb) as well as the same strain used by Krypotou et al. (2023) (*B. thetaiotaomicron* – Bth). Our results show that all chimeric proteins formed liquid droplets *in vitro*, similarly to AK390 (Figure 3E). This effect was stimulated by total RNA of *M. smegmatis*, except in the case of chimeric proteins that contain the IDR from Mab or Mhe (Figure 4F). Notably, the number of droplets significantly increased in the presence of RNA for all chimeras (Figure 4F). We also observed an increase in the number of large droplets for the IDR alone and for three of the four chimeric proteins, although the increase was only statistically significant for the chimera with the Mtb IDR (Figure 4G).

Altogether, these findings show that (i) _Msm_Rho undergoes LLPS *in vitro* due to the IDR, (ii) IDRs from other mycobacterial Rho proteins are sufficient to promote LLPS, and (iii) LLPS formation is promoted by RNA.

### 2.3 CRISPRi system to repress the native copy of _*Msm*_*rho*

We next investigated LLPS of _Msm_Rho *in vivo*. For these experiments, _Msm_Rho requires a fluorescent tag inside the *M. smegmatis* cells to allow the observation of condensates using confocal microscopy. Mutants with deletions of the Rho domains are also necessary to confirm the findings *in vitro*. We chose to repress the native copy of _*Msm*_*rho* in the wild-type *M. smegmatis* to facilitate expression of a second, tagged mutant copy of Rho.

CRISPR interference (CRISPRi) is a method to specifically repress the expression of target genes (Ghavami and Pandi 2021). We employed CRISPRi to knock down _*Msm*_*rho* expression using a single guide RNA (sgRNA) that targets the 5’ untranslated region (5’-UTR) of the gene. Applying a sgRNA design tool dedicated to Mycobacteria (Pebble tool: https://pebble.rockefeller.edu/tools/sgrna-design/), three sgRNAs for the _*Msm*_*rho* 5’ UTR were tested (Table S4 and Figure S4). In *M. smegmatis, rho* is an essential gene (MSRdb: Mycobacterial Systems Resource); therefore, *rho knockdown* is expected to lead to a reduction in growth rate. Our results showed substantial growth inhibition of *M. smegmatis* expressing each of two of the three sgRNAs in the presence of the anhydrotetracycline (aTc) inducer when compared to a control strain (Figure 5A). Similar results were obtained with the sgRNA positive controls that target expression of genes encoding DNA polymerase β, a hypothetical essential mycobacterial protein (MSMEG_0317), and Rho (targeting the protein-coding sequence) (Judd et al. 2021) (Figure 5A).

**Figure 5:**
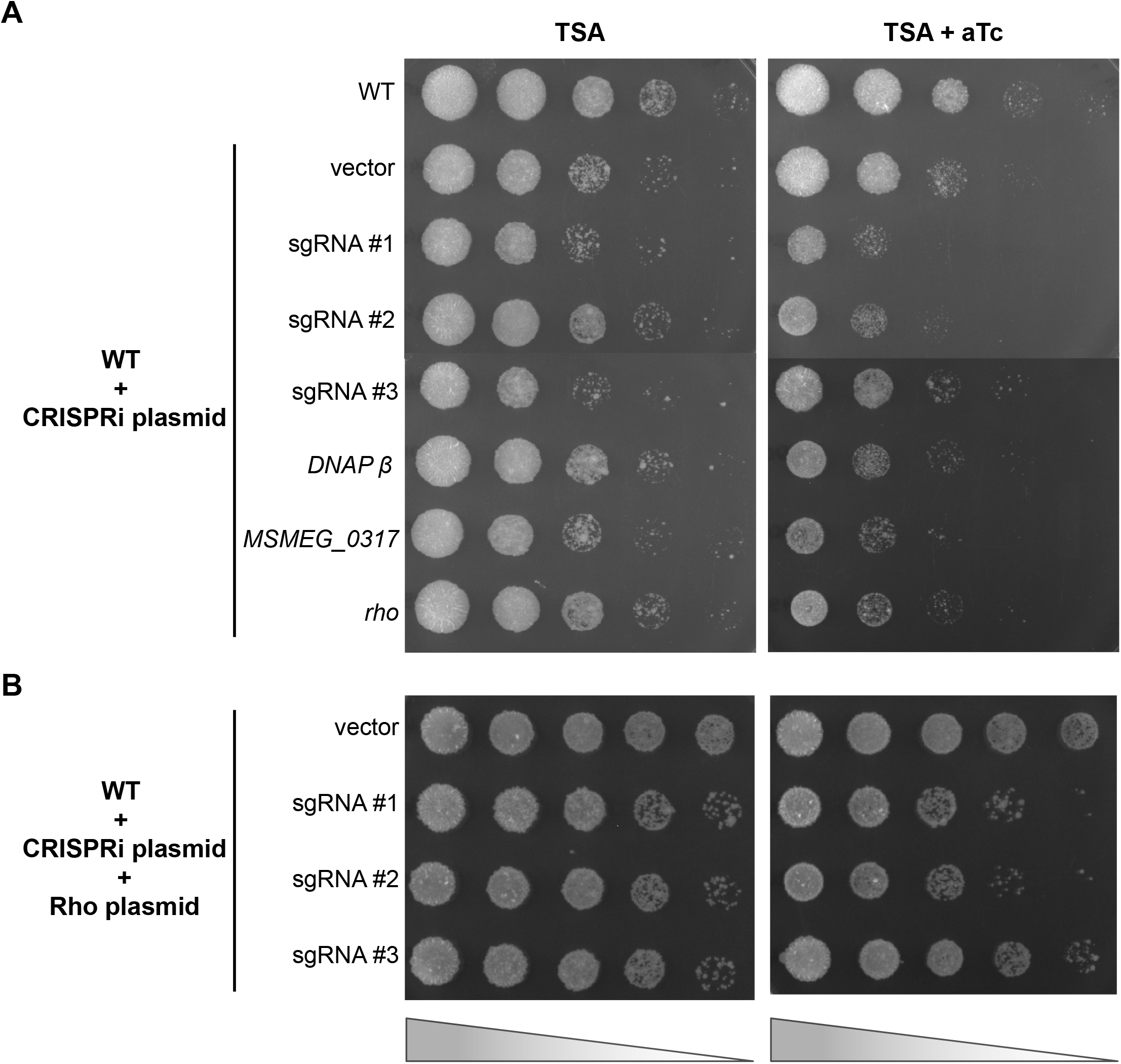
Substantial inhibition and restoration of the growth of *M. smegmatis* strains containing CRISPRi sgRNAs. (A) Wild-type strain harbouring the CRISPRi system with different sgRNAs for the promoter region of *rho* and three positive controls (*DNAβ, MSMEG_0317*, and *rho*). (B) Wild-type strain containing the CRISPRi plasmid and a second copy of *rho*. Strains with pJR962 derivatives expressing sgRNAs were grown to the stationary phase in TSBT in the absence of aTc. The cells were serially diluted 10-fold, and 5 µL of each dilution was spotted onto TSA without (left) or with (right) aTc, which induces expression of the CRISPRi system. Positive controls were designed by Judd et al. (2021).

Having obtained *M. smegmatis* strains with reduced expression of the native copy of *rho*, we introduced an integrative plasmid with a second copy of the *rho* gene. Growth of these cells was partially restored when compared to strains lacking the second copy of *rho* (Figure 5B).

Based on these results, we chose the cells containing the CRISPRi system with sgRNA #1 for further experiments.

### 2.4 IDR of _Msm_Rho plays a role in acid stress tolerance

The formation of membrane-less organelles by LLPS depends on pH, temperature, salt type, and salt concentration (Alberti et al. 2019). In this context, the optimal growth conditions of a bacterium might not be favorable to drive the phase separation of a protein inside the cells. Acting against stress is one of the possible outcomes of proteins that undergo phase separation in bacteria (Azaldegui et al. 2021; Gao et al. 2021). Therefore, we evaluated the effects of several stress conditions on the growth of the *M. smegmatis rho knockdown* mutants. We wanted to identify a potential scenario where the cultures with the wild-type Rho have increased fitness compared to the mutants with deletions in the Rho domains.

From the different conditions tested by Gebhard et al. (2008), all cultures grew well in the control (TSAT) and 4-hour exposure to ethanol, water, and basic pH (Figure S5A). Conversely, we observed differences in the growth of cultures exposed to high or low temperatures, high salt concentration, acid pH, and hydrogen peroxide treatment (Figure S5B).

The growth of the mycobacterial mutants seemed to be more affected by low pH. To capture the impact of acid stress on the mutant cultures, viable cell counts were determined, and a linear model with interaction terms was derived (Table 1). Although there was a significant variation in the values (Table S6), the model captured the negative impact on acid stress in the culture containing only the RNA-binding and the Rho ATPase domains (ΔIDR). In contrast, the effect of acid stress was not significant in all the other cultures. These results suggest that the IDR of _Msm_Rho is involved in the tolerance of *M. smegmatis* to acid stress.

### 2.5 _Msm_Rho undergoes phase separation *in vivo* after acid stress

In addition to counting the number of viable cells after exposure or not to pH 4 for four hours, we examined the subcellular localization of Dendra-tagged Rho and its derivatives (Figure 6). Confocal microscopy of the live cells revealed that for all constructs, Rho was regularly distributed along the cellular medial axis when cells were grown in the control medium (TSBT). This result is indicated by the constant yellow/orange (high-intensity fluorescence) displayed in the demographs. After acid stress, this same pattern was only observed for cells with Rho that lacks the IDR (Figure 6B). By contrast, cells with wild-type Rho (Figure 6A), or Rho lacking the initial (Figure 6C) or insertion (Figure 6D) domains individually showed patterns of multiple dispersed foci along the cells when exposed to acid stress. This observation is represented by the black/blue vertical slices with yellow/orange stripes in the demographs.

**Figure 6:**
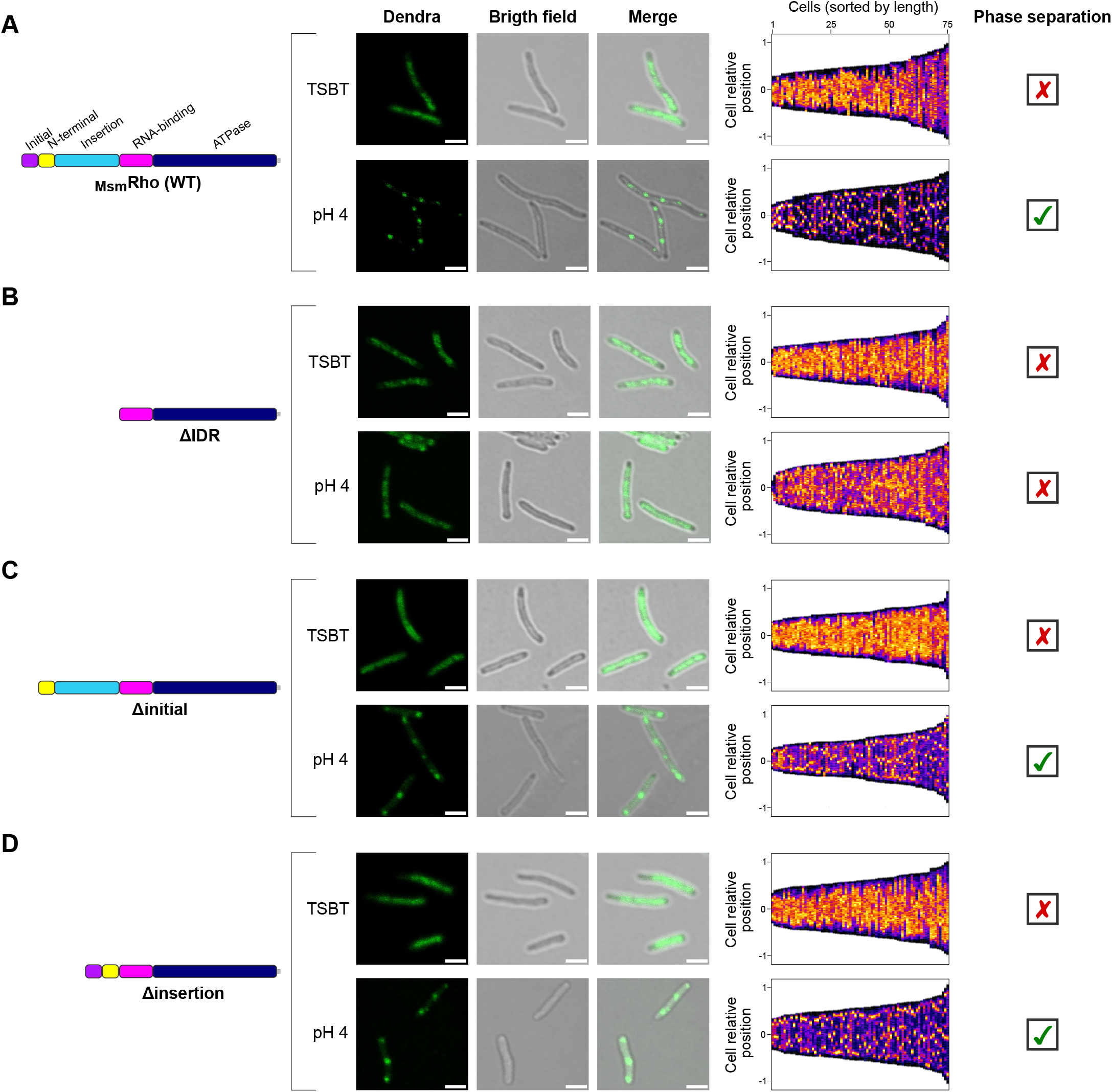
Additional regions _Msm_Rho mediate the formation of condensates *in vivo* after exposure to acid pH. Localization patterns for _Msm_Rho and its derivatives after 4-hour exposure to TSBT or pH 4 (citrate/phosphate buffer): (A) Wild-type Rho; (B) ΔIDR; (C) Δinitial; (D) Δinsertion. All cultures were supplemented with aTc to repress the native copy of *rho* through the CRISPRi system. Demographs on the right display the normalized Dendra intensity as heatmaps along the medial axis of single cells (n = 75) which are sorted by length. All analyses were carried out from three independent experiments, and representative images are shown. All scale bars, 2 µm.

Collectively, these results suggest that _Msm_Rho exhibits LLPS *in vivo* following a harsh challenge (acid stress), but not in the absence of stress. Moreover, phase separation in _Msm_Rho is dependent on the presence of the Rho additional regions.

## 3 DISCUSSION

Rho-dependent transcription termination is the primary mechanism for stopping transcription in Mycobacteria (Ahmad et al. 2023; D’Halluin et al. 2023). In fact, Rho in *M. tuberculosis* is not only crucial to preserve the integrity of transcription but also to establish and sustain infection in a mouse model (Botella et al. 2017). Mycobacterial Rho differs from *E. coli* Rho due to the presence of additional regions at the N-terminal region. Our data indicate that these additional regions promote the formation of Rho condensates through LLPS. Furthermore, the ability of Rho to form condensates may impact the ability of *M. smegmatis* to resist acid stress.

The three main domains (N-terminal, RNA-binding, and ATPase) are well conserved in mycobacteria (Figures S3 and S4), which agrees with the hypothesis of a common ancient origin of Rho in Bacteria (D’Heygère et al. 2013; Moreira et al. 2024). The predominant small insertion in the RNP3 of the RNA-binding domain has been reported before in Actinomycetota (*Micrococcus luteus*) and has been suggested to aid the binding of Rho to high GC transcripts (Nowatzke et al. 1997). Also abundant in Mycobacteria are the initial and insertion regions (Figure 2). All initial regions start with the same TDL motif. We speculate that threonine, aspartate, and leucine contribute to the protein’s stability based on the chemical and structural properties of these amino acids (Chou and Fasman 1973; Kisselev et al. 2000; Roesgaard et al. 2022). In the insertion regions, the additional arginines of the RxR motif may enhance binding to RNA. The RxR motif may also contribute to LLPS, since as has been shown for arginine-rich motifs in eukaryotes (Wang et al. 2018; Ottoz and Berchowitz 2020). The EDDV motif may function as an extension of the RNA-binding domain, given its location adjacent to this domain. By contrast, the NDQG motif is more divergent, and it could potentially be involved in conformational changes of Rho. Notably, asparagine and/or glutamine residues are enriched in domains of prion-forming proteins that assemble as amyloid aggregates (Sant’Anna et al. 2016; Shattuck et al. 2017).

Our data indicate that _Msm_Rho can undergo LLPS *in vitro* and *in vivo* following acid stress (Figure 6). Unlike LLPS of _Bth_Rho, which requires RNA, mycobacterial Rho can undergo LLPS in the absence of RNA, although RNA stimulates condensate formation. Consistent with computational predictions, the IDR is necessary and sufficient for LLPS. We observed differences in the growth of *M. smegmatis* with different Rho mutants following acid stress. These growth differences correlate with the ability of the Rho mutants to undergo LLPS *in vivo*, suggesting that LLPS contributes to cell fitness during acid stress. The fact that _Msm_Rho undergoes phase separation only after stress might be explained by the availability of RNA. In an acidic condition (pH 4), ribosomes are downregulated in *M. tuberculosis* and *M. smegmatis* (Choudhary et al. 2022; Cossu et al. 2013; Roxas and Li 2009). We speculate that there is more exposed mRNA available where the significant motifs (RxR and EDDV) of the Rho additional regions can bind. Upon binding to mRNA, _Msm_Rho assembles into condensates (Figure 7). By concentrating inside the droplets, _Msm_Rho might terminate the transcription of distinct transcripts that allow the tolerance to acid pH. To confirm this hypothesis, additional experiments of mapping of RNA termini (e.g., Term-seq) are necessary.

**Figure 7:**
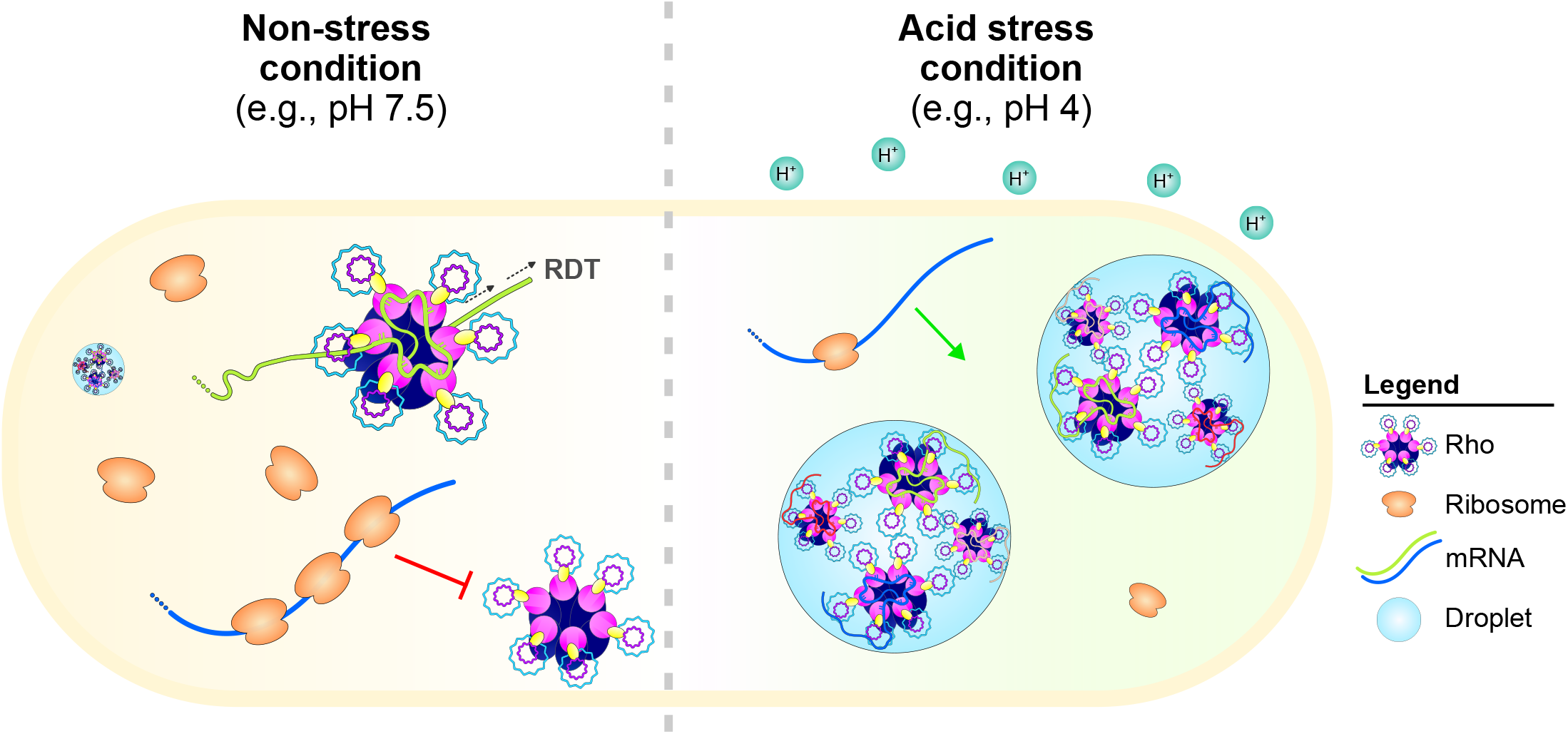
Proposed model for the assembly of _Msm_Rho into condensates during acid stress. The high ribosome occupancy of newly made transcripts blocks Rho binding to the rut sites and Rho-dependent termination (RDTT) occurs in levels that are not sufficient to promote LLPS in a non-stress condition (e.g., growth in TSBT with pH 7.5 – left part of the figure), although some small droplets might be formed. Ribosomes are downregulated during acid stress (e.g., pH 4 – right part of the figure) which increases the exposure of mRNA stimulating Rho binding and assembling into LLPS that occurs due to the presence of additional regions that are intrinsically disordered. In this scenario, Rho is concentrated within the condensates terminating different transcripts which allows acid tolerance.

In conclusion, _Msm_Rho undergoes phase separation, which may modulate acid tolerance. This process is likely to be stimulated by RNA and occurs due to the presence of additional protein regions that are likely intrinsically disordered and contain specific sequence motifs. Moreover, our results suggest that LLPS of Rho might be widespread in Mycobacteria, given the common initial and insertion regions with compositional bias and our *in vitro* assays that confirmed droplet formation of Rho containing additional regions of distinct mycobacterial genera. Our study has uncovered additional properties of Rho that are distinct from those of *E. coli*. Disruption of the phase separation of Rho might represent an interesting approach for the development of novel antimycobacterial therapies.

## 4 EXPERIMENTAL PROCEDURES

### 4.1 Bioinformatics analyses of mycobacterial Rho factors

#### 4.1.1 Mycobacterial Rho dataset

The annotations of the complete reference genomes from the Mycobacteriaceae family (n = 118) were downloaded from the NCBI database in June 2023 (Table S1). Rho sequences were identified by hmmsearch (Eddy 2011) as previously described (Moreira et al. 2024).

#### 4.1.2 Sequence alignment and logo

Multiple sequence alignment was performed using MAFFT (v7.394) (Katoh and Standley 2013) with default parameters and trimmed by trimAl (v1.4.rev220) (Capella-Gutiérrez et al. 2009) with a gap threshold of 0.05. The alignment was visualized on Geneious Prime software (v 2022.2.2) (http://www.geneious.com/) and color-coded according to the polarity scheme. The conservation pattern of the alignment was represented by a logo generated with the WebLogo 3 tool (https://weblogo.threeplusone.com) (Crooks et al. 2004) and color-coded based on amino acid chemistry.

#### 4.1.3 Bioinformatics predictions

The intrinsic disorder propensity was predicted with flDPnn (putative function- and linker-based Disorder Prediction using deep neural network) (http://biomine.cs.vcu.edu/servers/flDPnn/) (Hu et al. 2021). Spontaneous liquid–liquid phase separation and droplet-promoting regions were predicted using FuzDrop (https://fuzdrop.bio.unipd.it/) (Hatos et al. 2022). The scores of these programs were plotted against the Rho sequences with values between 0 and 1. Scores above 0.3 and 0.6 indicate IDRs and the ability to promote LLPS, respectively. The 3D structures were retrieved as PDB files from the AlphaFold Protein Structure Database (https://alphafold.ebi.ac.uk) (Varadi et al. 2022), and the domains were colored on PyMOL (v2.5.2) (Schrödinger, LLC).

#### 4.1.4 Motif discovery

Motif discovery analyses were conducted with MEME (Multiple Em for Motif Elicitation) Suite (v5.5.2) (Bailey and Elkan 1994) and MAST (v5.5.2) (Motif Alignment and Search Tool) (Bailey and Gribskov 1998) on mycobacterial Rho sequences. The detection of significant motifs (5-12 amino acids) was performed with MEME using a set of typical Rho sequences in the background model. The confirmation and location identification of these motifs were done with MAST.

#### 4.1.5 Phylogenetic reconstruction of Mycobacterial species

The phylogenetic tree showing the distribution of the significant motifs in the N-termini portion of the mycobacterial Rho sequences was generated with PhyloT (https://phylot.biobyte.de) based on NCBI taxonomy (Schoch et al. 2020). The motifs were annotated on the tree using iTOL (v6) (https://itol.embl.de/) (Letunic and Bork 2021).

### 4.2 Bacterial strains and growth conditions

Supplementary Table S2 describes all bacterial strains used in this study. *M. smegmatis* wild-type strain mc^2^155 and its derivatives were grown in tryptic soy broth (BD Bacto™ Tryptic Soy Broth, 211825) supplemented with 0.05% Tween 80 (AJAX FineChem, AJA2510-500) (TSBT) or on TSA plates. *E. coli* DH5α and BL21 (DE3) cultures were grown in Luria-Bertani (LB) medium. All liquid cultures were incubated overnight at 37 °C with shaking (225 rpm). All agar plates were incubated at 37 °C overnight (*E. coli* plates) or for three days (*M. smegmatis* plates). When necessary, antibiotic selection was used at the following concentrations: apramycin (12.5 µg/ml on agar, 10 µg/ml in broth), kanamycin (50 µg/ml and 10 µg/ml, respectively), and ampicillin (100 µg/ml).

### 4.3 Construction of recombinant strains

Supplementary Tables S3, S4, and S5 list all plasmids and oligonucleotides used in this study. Plasmids were constructed with NEBuilder® Hifi Assembly (NEB, E2621S) and extracted with QIAprep® Spin Miniprep Kit (QIAGEN, 27104) following the manufacturer’s instructions. Domain deletions were generated with Q5® Site-Directed Mutagenesis Kit (NEB, E0554S). The sequences of all constructed plasmids were verified by DNA Sanger sequencing.

Transformation of *E. coli* competent cells was achieved with a traditional heat-shock transformation. Preparation of *M. smegmatis* electrocompetent cells and electroporation were performed as described by van Kessel and Hatfull (2008). Details about the generation of all plasmids are presented in the following three sections.

#### 4.3.1 Construction of *E. coli* BL21 strains for expression and purification of Rho proteins

To test *in vitro* phase separation, a series of pET28a vectors was generated with the _Msm_Rho sequence codon optimized for *E. coli*. A construct containing genes coding for a His-tag followed by the _m_NeonGreen, a linker, and Rho cloned into the pET28a backbone was synthesized by Twist Bioscience. In addition, the optimized sequences of the N-terminal and the additional regions of different Mycobacteria species were synthesized: *M. abscessus* (Mab), *M. heraklionensis* (Mhe), *M. koreensis* (Mko), and *M. tuberculosis* (Mtb) as well as the same strain used by Krypotou et al. (2023) (*B. thetaiotaomicron* - Bth). These individual constructs were cloned into the pET28a vector before the RNA-binding domain of _*Msm*_*rho*.

Two types of deletions of the full-length _Msm_Rho in the _m_NeonGreen pET28a vector were generated to create constructs as follows: (i) ΔIDR: deletion of the first 855 bp of _*Msm*_*rho*; (ii) IDR: deletion of the last 1,140 bases of _*Msm*_*rho*. All resulting plasmids were transformed into *E. coli* BL21 (DE3).

#### 4.3.2 CRISPRi clones targeting the native copy of _Msm_Rho

To silence the expression of the native Rho, allowing only the effects of our constructs, we employed the CRISPR interference (CRISPRi) system with a single guide RNA (sgRNA) targeting the promoter region of *rho*. The idea was to preserve the expression of the second copy of Rho in the plasmids, which have a different promoter region. The sgRNA oligos were designed using the Pebble tool (https://pebble.rockefeller.edu/tools/sgrna-design/), which is specific for Mycobacteria. The three sgRNA target sequences with the highest predicted strengths were selected, and the sgRNA oligos were sent for synthesis.

The CRISPRi protocol to repress _Msm_Rho was performed following Wong and Rock (2021). The sgRNA oligos were cloned into the BsmBI-digested CRISPRi backbone (plJR962 - Addgene plasmid #115162). The CRISPRi plasmids were transformed into *E. coli* DH5⍰, and the colonies were selected with kanamycin. The confirmed vectors were transformed into electrocompetent *M. smegmatis* cells by electroporation. To validate the knockdown phenotype, the transformant cells were plated on TSA with or without 100 ng/mL anhydrotetracycline (aTc).

#### 4.3.3 Construction of *M. smegmatis* strains with a second copy of Rho

The vectors to test the *in vivo* separation of _Msm_Rho and its derivatives containing deletions of the domains were created based on the Rho Dendra plasmid (Dendra-MSMEG-4954) generated by Judd et al. (2021). First, the TetR operator site in the Rho Dendra plasmid was deleted, leading to the original constitutive mycobacterial promoter Psmyc (Ehrt et al. 2005). From this new plasmid, the new promoter, the *rho*, and the *dendra2* genes were amplified. The amplification products were cloned into the pYL180 vector, which has the apramycin resistance, and the Tweety integration site is compatible with the CRISPRi vectors. The final pYL281 vector had *dendra2* at the N-termini position of Rho. An untagged vector (pYL181) without *dendra2* was also generated.

To assess the role of the initial and/or the insertion regions of _Msm_Rho in surviving stress and phase separation, a series of deletions/replacements of the full-length _Msm_Rho was performed to generate new functional Rho plasmids. Therefore, the constructs contain at least the RNA-binding and the ATPase domains without both or one of the Rho additional regions. The _Msm_Rho mutants were generated as follows: (i) ΔIDR: deletion of the first 855 bp of _*Msm*_*rho*; (ii) Δinitial: replacing the first 102 bp of _*Msm*_*rho* with the first 12 bp from E. coli rho (_*Ec*_*rho*); (iii) Δinsertion: replacing the region comprehending the bases 211 to 855 of _*Msm*_*rho*. The Rho plasmids were transformed into *E. coli* DH5⍰ and the colonies were selected with apramycin. The confirmed vectors were transformed into electrocompetent CRISPRi *M. smegmatis* cells by electroporation.

### 4.4 Expression and purification of Rho proteins

The purification of recombinant Rho proteins was performed following Krypotou et al. (2023) with some modifications. Cultures were grown at 37 °C in 200 mL of LB with 50 µg/mL kanamycin until reaching an OD_600 nm_ of 0.5. Next, 0.2 mM IPTG was added, and the cultures were incubated at 30 °C for 6 hours. Cells were centrifuged and the pellet was resuspended in 5 mL of lysis buffer [50 mM Tris HCl pH 7.5, 1 M NaCl, 5% (vol/vol) glycerol, and 0.1 mM DTT, supplemented with cOmplete^TM^, Mini, EDTA-free Protease Inhibitors cocktail (Roche, 11836170001)] following by sonication at amplitude: 35%, pulse: 1.5 sec on/ 0.5 sec off and total time of 3 min. The lysate was centrifuged at 3,200 x g at 4 °C for 15 min. The supernatant was applied onto 0.5 mL HIS-Select® Nickel Affinity Gel (Merck, P6611) previously washed with lysis buffer and loaded on Poly-Prep® Chromatography Columns (Bio-Rad, 7311550) for gravity flow chromatography. The nickel resin was washed twice with 1 mL lysis buffer, followed by 1 mL lysis + 10 mM imidazole. The final elution was performed with 0.5 mL lysis buffer + 200 mM imidazole. The protein concentration was quantified by NanoDrop. The eluted proteins were concentrated by ultrafiltration (Amicon-Millipore, UFC503024), and the buffer was exchanged into a storage buffer [100 mM Tris pH 8, 500 mM KCl, 50% (v/v) glycerol, 0.1 mM DTT].

The protein purity was analyzed with SDS-page which the samples (5 µL) were diluted in 2X Laemmli sample buffer (Bio-Rad, 1610737) supplemented with 0.1 mM DTT, heated at 95 °C for 5 min, and separated on 4–20% Mini-PROTEAN® TGX™ Precast Protein Gels (Bio-Rad, 4561094). The gel was run in 1X diluted from the 10X Tris/Glycine/SDS buffer (Bio-Rad, 1610732) for 90 min at 95 V and stained with Coomassie brilliant blue R-250 (Figure S6). Protein samples were aliquoted and stored at −80 °C.

### 4.5 Total RNA extraction from *M. smegmatis*

Total RNA from *M. smegmatis* wild-type strain mc^2^155 was extracted using TRIzol (Invitrogen, 15596026) with mechanical lysis of the cells. Briefly, cells were harvested after overnight growth in TSBT and resuspended in 1 mL TRIzol. Next, the mixture was homogenized carefully by pipetting, incubated at room temperature for five minutes, and transferred to the Lysing Matrix E tubes (MP Biomedicals, 116914050-CF). The mechanical lysis was performed with BeadBug™ 6 microtube homogenizer (Merck, Z742684) (power: 280/ 2 cycles of 1 min). Following centrifugation (12,000 x g/15 min/ 4 °C), chloroform was added to the supernatant, mixed carefully by inversion, and the tubes were centrifuged again (12,000 x g/15 min/ 4 °C). The colorless upper aqueous phase was transferred to a fresh tube, and RNA was precipitated by adding isopropanol, mixing carefully by inversion, and incubating the samples at −20 °C overnight. Next, the tubes were centrifuged (12,000 x g/15 min/ 4 °C) and the RNA pellets were washed with 75% ethanol, followed by the last centrifugation step (7,500 x g/5 min/ 4 °C). After air-drying, the RNA pellets were resuspended in 20 µL of RNase-free water (Rio et al. 2010).

### 4.6 *In vitro* phase separation assays

The buffer in the purified proteins was switched to a high salt buffer [50 mM Tris pH 7.5, 500 mM KCl, 2.5% (v/v) glycerol, 0.1 mM DTT] using ultrafiltration. Depending on the desired salt and protein concentrations, the samples were diluted in a buffer lacking salt [50 mM Tris pH 7.5, 2.5% (v/v) glycerol, 0.1 mM DTT]. Proteins (2 µL) were pipetted on the functionalized glass slides and imaged immediately.

Glass slides (Ibidi, 81826) were functionalized at room temperature as follows: (i) incubation with 0.5% Hellmanex III (Merk, Z805939) for one hour and washing three times with MilliQ water; (ii) incubation with 10 M NaOH for 15 min and washing three times with MilliQ water; (iii) incubation with 0.1 mg/mL PLL(2)-g[3.5]-PEG(2) (SuSoS, Dubendorf, Switzerland) dissolved in 10 mM HEPES (pH 8.0) during two days and washing five times with MilliQ water; (iv) air drying the functionalized slides with compressed N_2_. Slides were stored at −20 °C until the time of use.

### 4.7 Stress condition experiments

Stress conditions experiments were performed as Gebhard et al. (2008) with modifications. Mycobacterial cultures were grown overnight in the TSBT medium, and the OD_600 nm_ was adjusted to 0.2 for all conditions. For temperature stress, cultures were incubated at 50 °C (heat-shock) and 18 °C (cold-shock). For pH stress, cells were harvested by centrifugation (8,000 g/ 3 min) and resuspended in citrate/phosphate buffer (150 mM, pH 4) (acid stress) or sodium borate buffer (13 mM, pH 9) (alkaline stress). For osmolarity stress, cells were harvested as previously described and resuspended in 0.5 M NaCl (hyperosmotic stress) or distilled water (hypoosmotic stress). The ethanol (5%) and oxidative (7.5 mM H_2_O_2_) stresses were performed by adding the compounds directly to the OD_600 nm_-adjusted cultures. All solutions were supplemented with 0.5% Tween 80 and 100 ng/mL aTc. Cultures were incubated under different stress conditions for 4 hours at 37 °C (or the stated temperature) with agitation (225 rpm). The only exception was the 2 hours of incubation for the oxidative stress. Then, the cultures were diluted until 1:10^-5^ and the dilutions were plated on TSA complemented with 100 ng/mL aTc. The counting of the viable cells before and after acid stress was performed with the drop plate method as described by Herigstad et al. (2001) from five replicate spots of 10 µL per dilution.

### 4.8 *In vivo* phase separation assays

*M. smegmatis* strains were grown overnight in TSBT supplemented with 100 ng/mL aTc. Cells were collected by centrifugation (8,000 g/ 3 min) and resuspended in the citrate/phosphate buffer (150 mM, pH 4) with 0.05% Tween 80 or TSBT, both added 100 ng/mL aTc. The cultures were incubated for 4 hours at 37 °C with agitation (225 rpm). Next, the cells were concentrated after centrifugation (8,000 g/ 3 min), and 5 µL were loaded onto a 35-mm glass-bottom dish (Cellvis, D35-20-1-N). Cell immobilization was achieved by 1.5% agarose pads placed on the top of the cell dish, covered by a coverslip.

### 4.9 Confocal microscopy

All microscopy images were acquired with an Olympus FV3000 inverted microscope under the 100x oil immersion objective lens. The _m_NeonGreen fluorescence of the purified Rho proteins was detected using the Dronpa-Green filter (excitation wavelength: 504 nm/emission wavelength: 518 nm). The Dendra2 fluorescence of the mycobacterial recombinant strains was detected with the DiO filter (excitation wavelength: 489 nm/emission wavelength: 506 nm).

### 4.10 Fluorescence Recovery After Photobleaching (FRAP) of _Msm_Rho droplets

Droplets were assembled with 5 µM of _Msm_Rho in 200 mM KCl buffer as described above for the *in vitro* phase separation. Samples were observed under a Zeiss LSM+ 900 with Airyscan 2 with a 63x oil immersion objective lens. A region within a droplet was selected, and the fluorescence signal was bleached with the 488-nm laser at 100% laser power for approximately 2 seconds. Time-lapse images were captured before and after the photobleaching at a rate of 1.5 seconds for about 3 minutes. The fluorescence recovery was calculated by normalizing the fluorescence intensity of the bleached area at each time point by the fluorescence intensity of a nearby, unbleached area over time.

### 4.11 Image analysis

Microscopy images were analyzed with Fiji-ImageJ (v2.14.0/1.54f). For the *in vitro* assays, the images of the fluorescent channel were selected, and an automatic threshold was set to convert the pixels in the images (droplets) to black and all the pixels above the threshold to white (background). Next, the “Analyse Particles” command in Fiji-ImageJ was used to identify the droplets with an area > 1 pixel^2^ and circularity of 1. When analyzing the effect of different KCl concentrations, all particles were detected without the circularity threshold. The intensity of the fluorescence in the cells was quantified using the MicrobeJ (v.5.13p) plugin (Ducret et al. 2016). Cell boundaries (n = 75) were identified with the medial axis algorithm in the bright field channel with the profile option and minimum cell length of 1 µm. Demographs of Dendra2 signal intensity across cell lengths after 4-hour incubation in TSBT or pH 4 buffer were built using the “Demograph” feature of MicrobeJ by plotting the medial intensity profiles of Dendra2 signals.

### 4.12 Statistical analysis

Using the “lm” function in R, a linear model with interaction terms was utilized to capture: (i) the variation in baseline colony-forming unity (CFU) across the different cultures; (ii) the varying baseline growth rates among these cultures; (iii) the overall impact of acid stress on growth rate; (iv) the differential effect of acid stress on growth rate across different cultures:

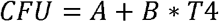

The constant term (A) is modelled as a function of the mutations of interest (wild-type, ΔIDR, Δinitial, and Δinsertion) to address the variation of baseline CFU (time = 0). When defining binary-indicator variables for each culture (C281-C284), we have:

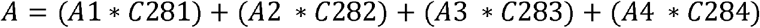

The coefficient (B) measures the CFU change relative to the baseline CFU and is used as a proxy for cell growth between the two time points (times 0 and 4 hours). It is modelled as a function of mutations of interest (C281-C284), and furthermore, each mutant responds to acidity (pH4) differently:

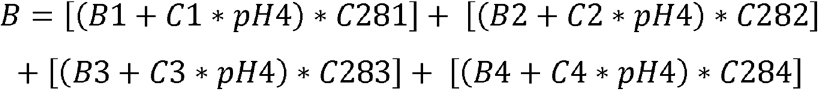

Each culture has its own base growth rate (B1-B4). When pH 4, the growth rate changes by different amounts, C1-C4, depending on culture types. The binary indicator (T4) refers to time = 4 hours.

## Author Contributions

**Sofia M. Moreira**: conceptualization, investigation, writing – original draft, methodology, validation, formal analysis, visualization. **Joseph T. Wade**: conceptualization, supervision, writing – review and editing. **Chris M. Brown:** conceptualization, supervision, writing – review and editing, funding acquisition, project administration, resources.

## Acknowledgments

C.M.B. was supported by a Marsden Grant (grant number: MFP-UOO1901) that also supported S.M.M. through a doctoral scholarship. JTW was supported by a grant from the National Institutes of Health (R35GM144328). We thank Dr. Eduardo Groisman for providing the *Bacteroides* strains and Rho constructs to serve as controls for the experiments. We also thank Dr. Amy Yewdall for her advice and assistance with the liquid-liquid phase separation experiments. We thank Dr. Te-yuan Chyou for developing the linear model.

## Data Availability Statement

The data that supports the findings of this study are available in the Supporting Information of this article.

---

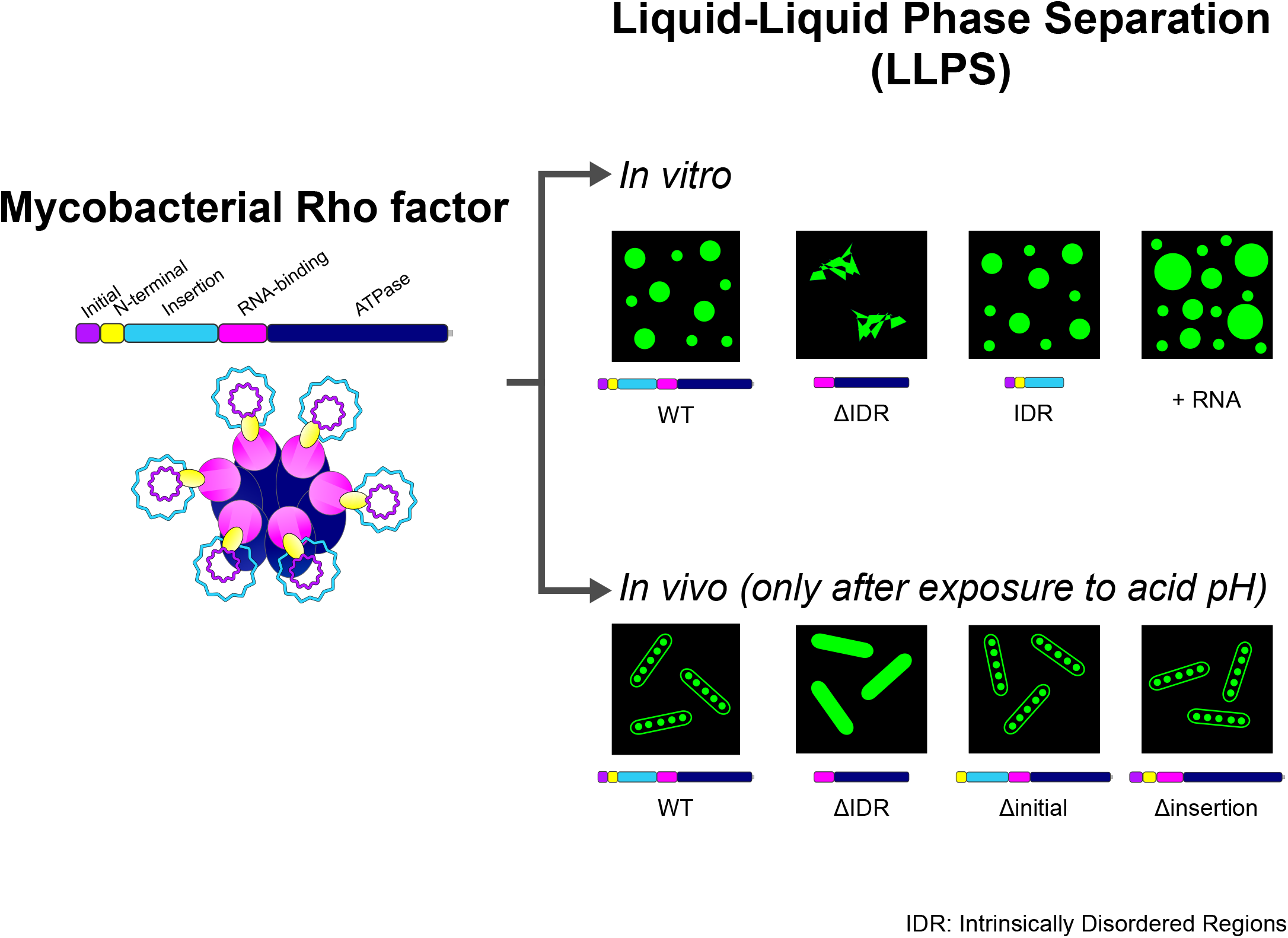

