## Supplementary Material for "Phase separation of mycobacterial Rho factor is associated with acid stress"

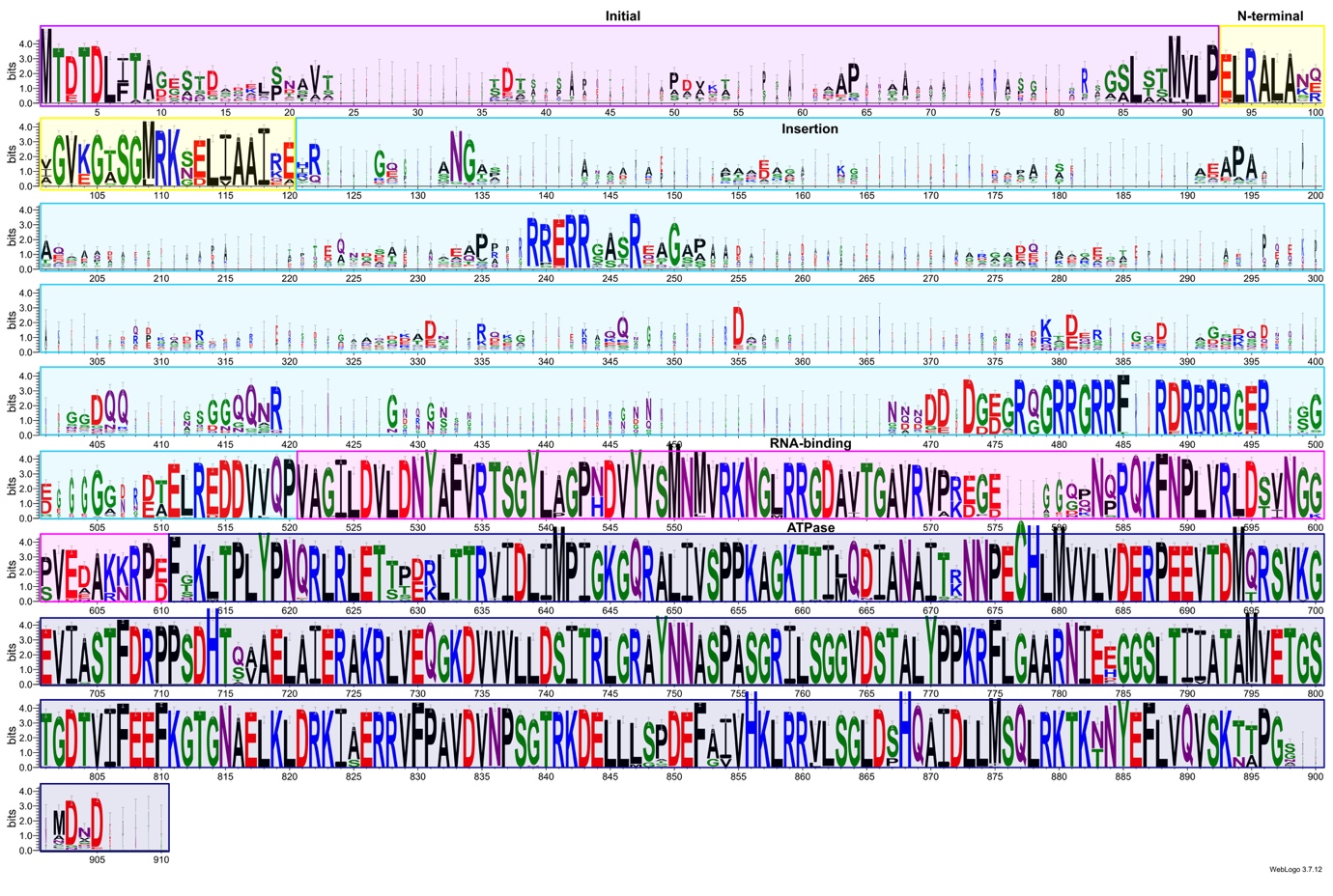

**Figure S1:** **Sequence logo of Mycobacterial Rho (n = 118)**. Rho domains are highlighted in different colours. Logo was generated from the sequence alignment on the WebLogo 3 server (https://weblogo.threeplusone.com).

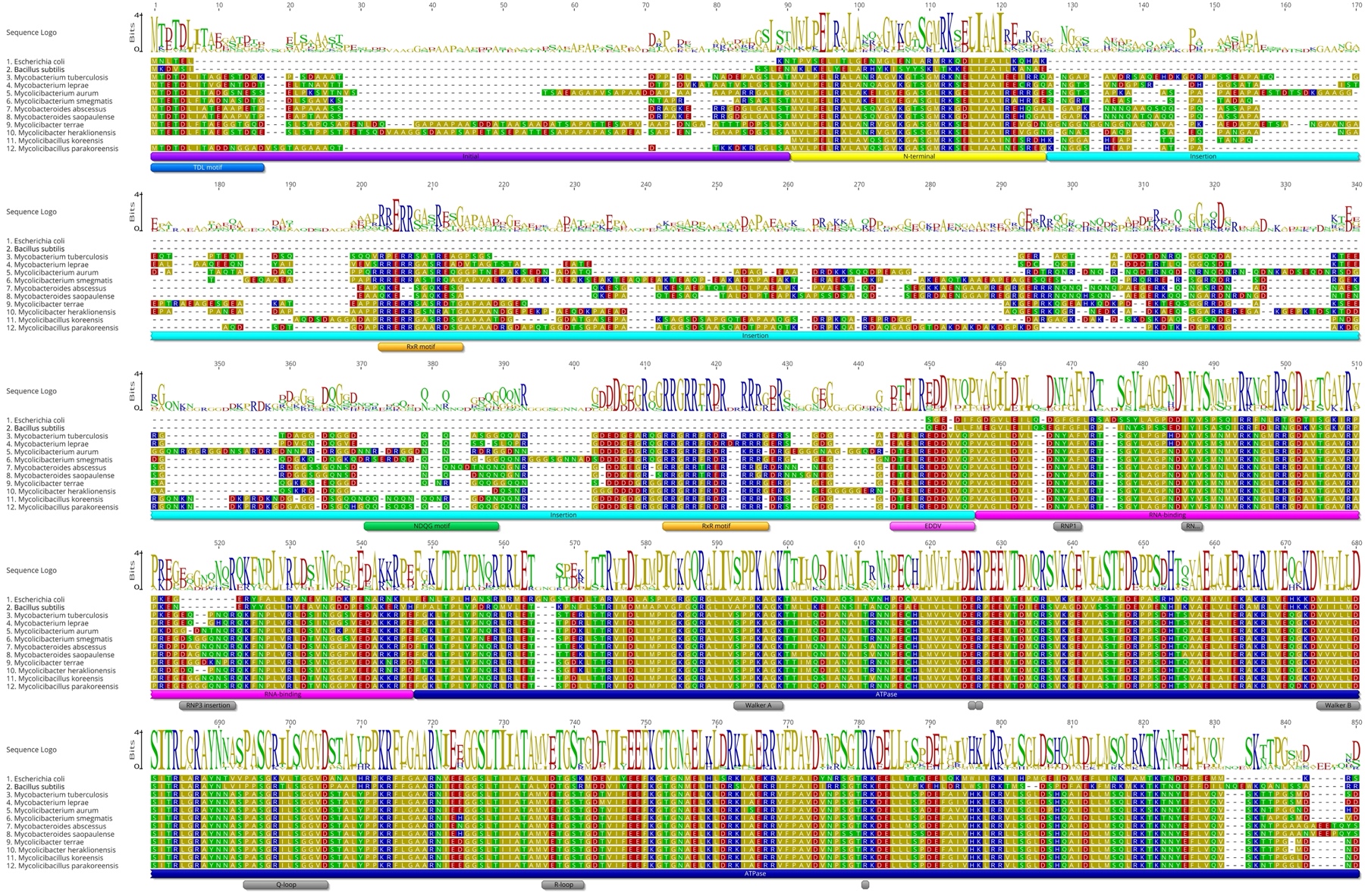

**Figure S2:** **Sequence alignment of Mycobacterial Rho (n = 10).** The sequences of _Ec_Rho and _Bs_Rho (*Bacillus subtilis*) were used as reference. Alignments were performed with MAFFT and coloured according to the polarity scheme in Geneious Prime.

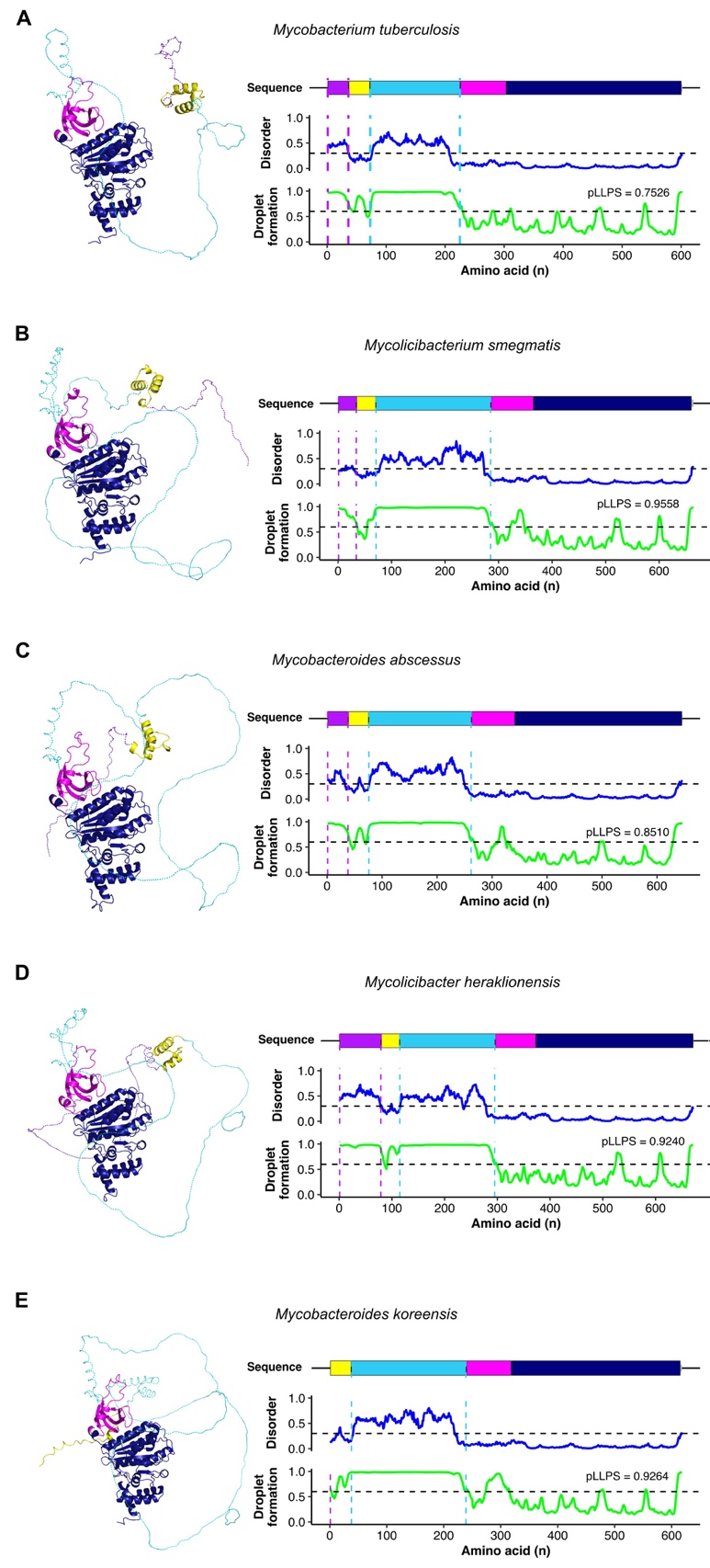

**Figure S3:** **Characterization of Mycobacterial Rho for disorder and droplet formation.** (A) *Mycobacterium tuberculosis* (AF-P9WHF3-F1). (B) *Mycolicibacterium smegmatis* (AF-I7G6F1-F1). (C) *Mycobacteroides abscessus* (AF-A0A829QAQ3-F1). (D) *Mycolicibacter heraklionensis* (AF-A0A1A2VQB2-F1). (E) *Mycobacteroides koreensis* (AF-A0A7I7S7N1-F1). 3D structures were extracted from the AlphaFold database and were coloured according to the different domains. The initial (purple) and insertion (light blue) regions were not confidently modelled probably due to their disordered nature. Predictions were performed by flDPnn (disorder) and FuzDrop (droplet formation). Abbreviation: pLLPS: Probability of forming a droplet state through liquid-liquid phase separation. Proteins with pLLPS ≥ 0.60 are droplet-drivers (can spontaneously undergo liquid-liquid phase separation).

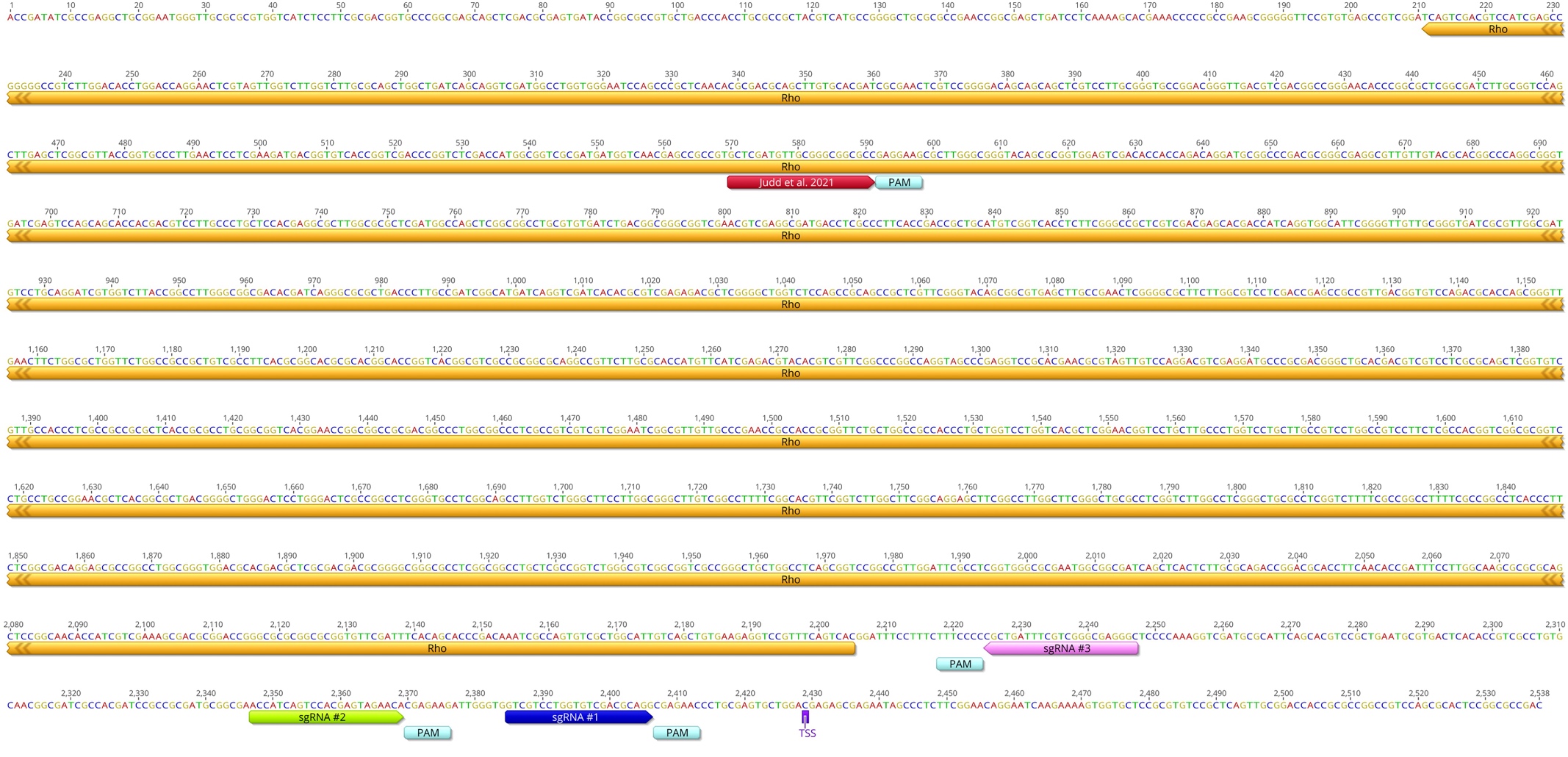

**Figure S4:** **Sequence of the *rho* gene in *M. smegmatis* mc^2^155 with the sgRNAs used in this study.** The sgRNA oligos (in blue, green, and pink) were designed with the Pebble tool (https://pebble.rockefeller.edu/tools/sgrna-design/). The *rho* sgRNA designed by Judd *et al.* (2021) was used as control. The Transcription Start Site (TSS) was retrieved from Martini *et al.* (2019).

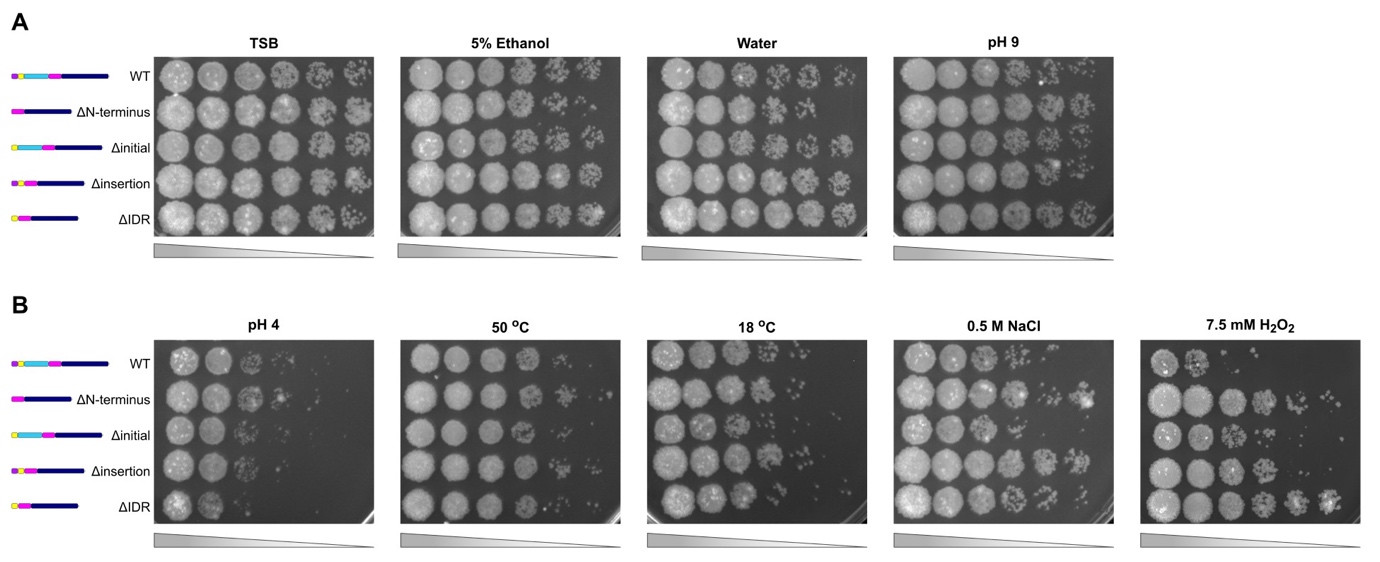

**Figure S5:** **Stress response of *M. smegmatis rho* knockdown strains harbouring a second plasmid with different deletions of the Rho domains.** (A) Conditions that did not affect the growth of the mutants. (B) Conditions that affected the growth of the mutants. Cultures were exposed to the different stress conditions for 4 hours (exception: 7.5 mM H_2_O_2_ for 2 hours) and 10-fold serial dilutions were plated on TSA. All solutions and media were supplemented with aTc to repress the native copy of *rho* through the CRISPRi system.

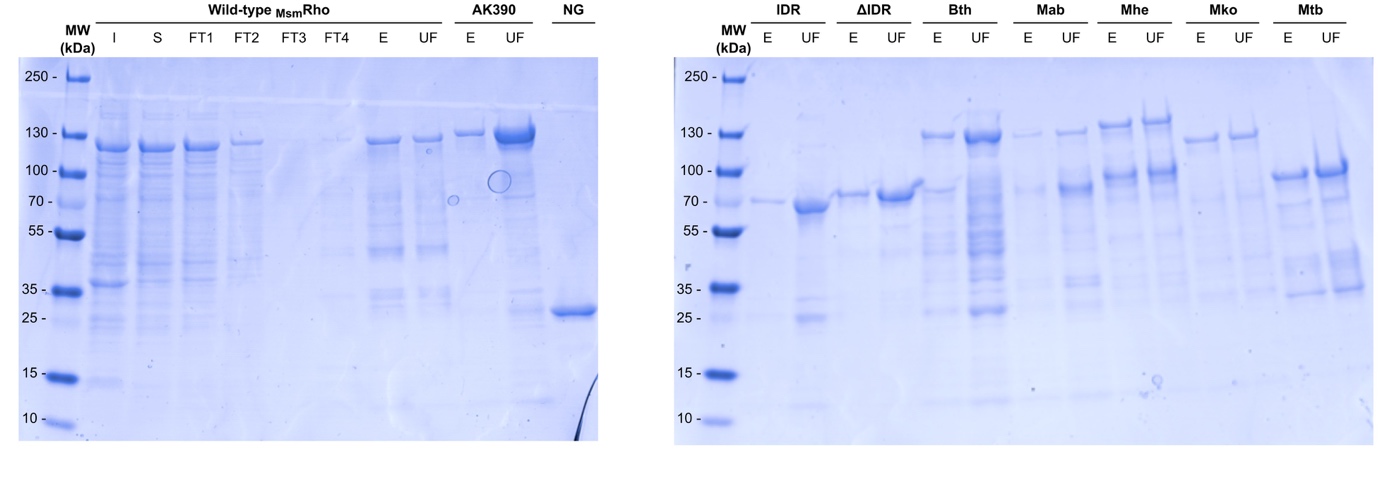

**Figure S6:** **Coomassie stained SDS-gels of the purified proteins used for the *in vitro* phase separation assays.** Fractions shown correspond to insoluble (I), soluble (S), first flowthrough (FT1), second flowthrough (FT2), third flowthrough (FT3), fourth flowthrough (FT4), eluted (E), and ultrafiltrate (UF) of the wild-type Rho (*Mycolicibacterium smegmatis*). Sample AK390 is the *Bacteroides thetaiotaomicron* Rho from (Krypotou *et al.*, 2023). All proteins have a His-tag and the _m_NeonGreen (NG) genes attached. Abbreviations for the fused proteins contained the IDR from different species and the RNA-binding and ATPase domains from *M. smegmatis*: Bth: *Bacteroides thetaiotaomicron*; Mab: *Mycobacteroides abscessus*; Mhe: *Mycolicibacter heraklionensis*; Mko: *Mycobacteroides koreensis*; Mtb: *Mycobacterium tuberculosis.*
